# Predicting protein functions using positive-unlabeled ranking with ontology-based priors

**DOI:** 10.1101/2024.01.28.577662

**Authors:** Fernando Zhapa-Camacho, Zhenwei Tang, Maxat Kulmanov, Robert Hoehndorf

## Abstract

Automated protein function prediction is a crucial and widely studied problem in bioinformatics. Computationally, protein function is a multilabel classification problem where only positive samples are defined and there is a large number of unlabeled annotations. Most existing methods rely on the assumption that the unlabeled set of protein function annotations are negatives, inducing the *false negative* issue, where potential positive samples are trained as negatives. We introduce a novel approach named PU-GO, wherein we address function prediction as a positive-unlabeled ranking problem. We apply empirical risk minimization, i.e., we minimize the classification risk of a classifier where class priors are obtained from the Gene Ontology hierarchical structure. We show that our approach is more robust than other state-of-the-art methods on similarity-based and time-based benchmark datasets. Data and code are available at https://github.com/bio-ontology-research-group/PU-GO.

## Introduction

Deciphering the functions of proteins is essential for unraveling the complexities of cellular pathways (Eisenberg *et al*., 2000), identifying potential drug targets (Schenone *et al*., 2013), and understanding diseases (Liu *et al*., 2015). In bioinformatics, protein function prediction emerges as a formidable challenge. With the rapid growth of biological data, including genomic and proteomic information, there is a pressing need for effective computational methods to predict protein functions accurately. Currently, the Uniprot Knowledge Base (UniprotKB) (Consortium, 2022) contains more than 250 million protein sequences and only few of them have experimental functional annotations. The Gene Ontology (GO) (Ashburner *et al*., 2000) provides structured information about protein functions and describes more than 50000 functions in three sub-ontologies: Molecular Function Ontology (MFO), Cellular Component Ontology (CCO) and Biological Process Ontology (BPO).

Despite substantial progress in bioinformatics, the functional annotations of proteins remain incomplete. A significant portion of the proteome lacks detailed functional characterization, hindering our comprehensive understanding of cellular processes. This incompleteness stems from the limitations of experimental techniques and the resource-intensive nature of functional assays. As a result, computational methods play a pivotal role in filling these knowledge gaps and providing predictions for unannotated or poorly characterized proteins.

In the pursuit of accurate protein function prediction, many existing methods adopt a binary classification learning framework, optimizing classifiers using unlabeled protein-function annotations as negative samples. This traditional approach, while effective in certain contexts, overlooks the nuances inherent in the protein function prediction landscape. Unlabeled samples might hide positive protein function annotations yet to be discovered.

UniprotKB regularly introduces new annotations for certain proteins; for example, from UniprotKB version 2023 03 to UniprotKB version 2023 05, there were 2,689 proteins that gained 4,362 functional annotations. Protein functional annotations can be propagated using the true-path rule, which results in 26,865 propagated annotations between both versions of UniprotKB. The oversimplified binary approach may lead to biased predictions and overlook potentially valuable information embedded in unlabeled proteins.

Positive unlabeled (PU) learning represents a paradigm shift in addressing these challenges. PU learning acknowledges the inherent uncertainty in the functional status of unlabeled protein function annotations and recognizes them as potential positives. In the PU learning realm, there are various approaches to handle unlabeled data (Bekker and Davis, 2020).

PU learning has been applied to different bioinformatics tasks (Li *et al*., 2021) such as disease gene predictions (Yang *et al*., 2012; Vasighizaker and Jalili, 2018; Stolfi et al., 2023), drug-target interaction prediction (Lan *et al*., 2016; Peng et al., 2017) as well as protein function prediction (Youngs *et al*., 2013; Song *et al*., 2021). There are two main strategies in which PU learning has been applied: negative extraction from the unlabeled data and probabilistic adaptation of a classifier (Li *et al*., 2021). Negative-extraction methods are a two-step process where a subset of *reliable negatives* are extracted from the unlabeled set and then a classifier is optimized with a conventional learning algorithm. Although this approach can show effectiveness across different bioinformatics tasks, the strategy of pre-selecting negatives can exclude important samples, producing inaccurate or biased classifiers.

Methods that adapt a classifier do not need to estimate a negative sample set *a priori*. Instead, the classifier is optimized with the whole dataset (positive and unlabeled) and estimation of positives/negatives from the unlabeled set are performed afterwards. These methods rely on the probabilistic formulation defined by Elkan and Noto (2008) for PU learning.

In the context of function prediction, most methods follow the negative samples extraction strategy (Zhao *et al*., 2008; Chen *et al*., 2010; Youngs *et al*., 2013), meaning that training is done with a fraction of the given data. On the other hand, methods that learn a classifier with PU data directly (Song *et al*., 2021) rely on optimization frameworks such as Majorization Minimization (Kenneth Lange and Yang, 2000) or Support Vector Machines (Cortes and Vapnik, 1995). However, in recent years, protein function prediction has been extensively addressed with emerging deep learning techniques(Kulmanov *et al*., 2017; Cao and Shen, 2021a; Yuan et al., 2023a; Wang et al., 2023).

We present PU-GO a method for predicting protein functions by optimizing a classifier under PU learning framework. Instead of pre-selecting negatives samples, PU-GO uses the classifier adaptation approach and minimizes classifications risks of positive and unlabeled samples (du Plessis *et al*., 2014). Our framework uses the ESM2 15B protein language model (Lin *et al*., 2022) to obtain high-dimensional feature vectors for protein sequences, which are used to optimize a multi-layer perceptron (MLP) classifier. Instead of enforcing the classifier to strictly discriminate between positive and negative samples, we use a ranking-based loss (Tang *et al*., 2022) to guide the classifier to rank positive samples higher than unlabeled ones. Furthermore, since protein function is a multi-label classification problem, we rely on the GO hierarchical structure to construct class priors for each GO function.

In this way, PU-GO aims to optimize a classifier in a more nuanced and accurate way for protein function prediction. This approach holds promise in enhancing the sensitivity and specificity of predictions, thereby contributing to a more comprehensive and reliable understanding of protein functions in complex biological systems. We show that PU-GO can outperform state-of-the-art protein function prediction methods in a similarity-based and time-based benchmark datasets.

## Materials and Methods

### Positive-Negative (PN) classification

Let **x** ∈ ℝ^*d*^ and *y* ∈ {−1, +1} be random variables with probability density function *p*(**x**, *y*) (du Plessis *et al*., 2014). Let *g* : ℝ^*d*^ → ℝ be an arbitrary decision function and *l* : ℝ → ℝ_+_ a loss function. The binary classifier *g* minimizes the risk:

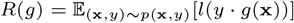

where 𝔼 is the expected value over *p*(**x**, *y*)

In standard binary classification, positive *P* and negative *N* data sets are given with distributions *p*_*P*_ (**x**) = *p*(*x*|*y* = +1) and *p*_*N*_ (**x**) = *p*(**x**|*y* = −1) (du Plessis *et al*., 2014). Given *π* = *p*(*y* = 1) as the prior for *P*, the risk *R*(*g*) can be expressed as:

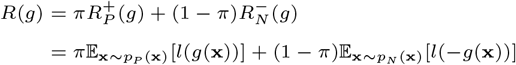

Assuming data from *P* and *N* are sampled independently, *R*(*g*) can be approximated by:

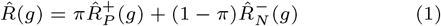

where 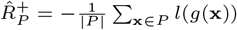 and 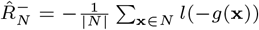

### Positive-Unlabeled (PU) classification

In PU classification, we assume the set *N* is empty and we are given an unlabeled data set *U* with marginal probability density function *p*(**x**). In this case, the risk 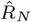 cannot be computed. However, we can express 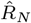using the following equality (Plessis *et al*., 2015):

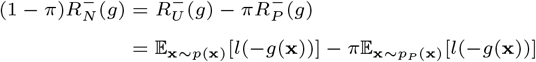

and Equation 1 becomes:

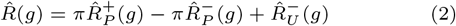

where 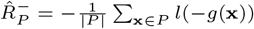 and 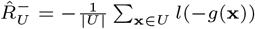. To avoid cases where *R*(*g*) can become negative, a non-negative estimator (Kiryo *et al*., 2017) is formulated as follows:

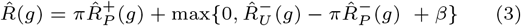

where 0 ≤ *β* ≤ *π*. Since *β* ≤ *π*, we construct it using a margin factor hyperparameter *γ*, such that *β* = *γπ*, with 0 ≤ *γ* ≤ 1.

### PU Learning for function prediction

In the context of function prediction, the feature space for **x** and functions *l* and *g* must be defined. We use the ESM2 15B (Lin *et al*., 2022) model to generate vectors for protein sequences that are consequently used as feature space **x**. The ESM2 15B model generates vectors of size 5120 that we refer to as *ESM2 vectors*.

We implement the classifier *g* as a multi-layer perceptron (MLP) that takes ESM2 vectors as inputs and returns values in ℝ^*k*^, where *k* is the number of classes. This classifier has shown to be effective in previous works (Kulmanov and Hoehndorf, 2022). The MLP network contains two layers of MLP blocks where the output of the second MLP block has residual connection to the first block. This representation is passed to the final classification. One MLP block performs the following operations:

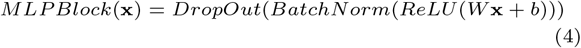

The input vector **x** of length 5120 represents ESM2 emedding and is reduced to 2048 by the first MLPBLock:

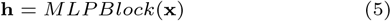

This representation is passed to the second MLPBlock with the input and output size of 2048 and added to itself using residual connection:

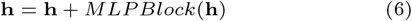

Finally, we pass this vector to a classification layer The output size of this layer is the same as the number of classes in each sub-ontology:

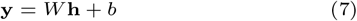

For PU learning, the loss function *l*(*x*) is:

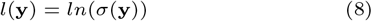

where *σ*(*x*) = 1*/*(1 + *e*^−*x*^) is the sigmoid function.

**Fig. 1.**
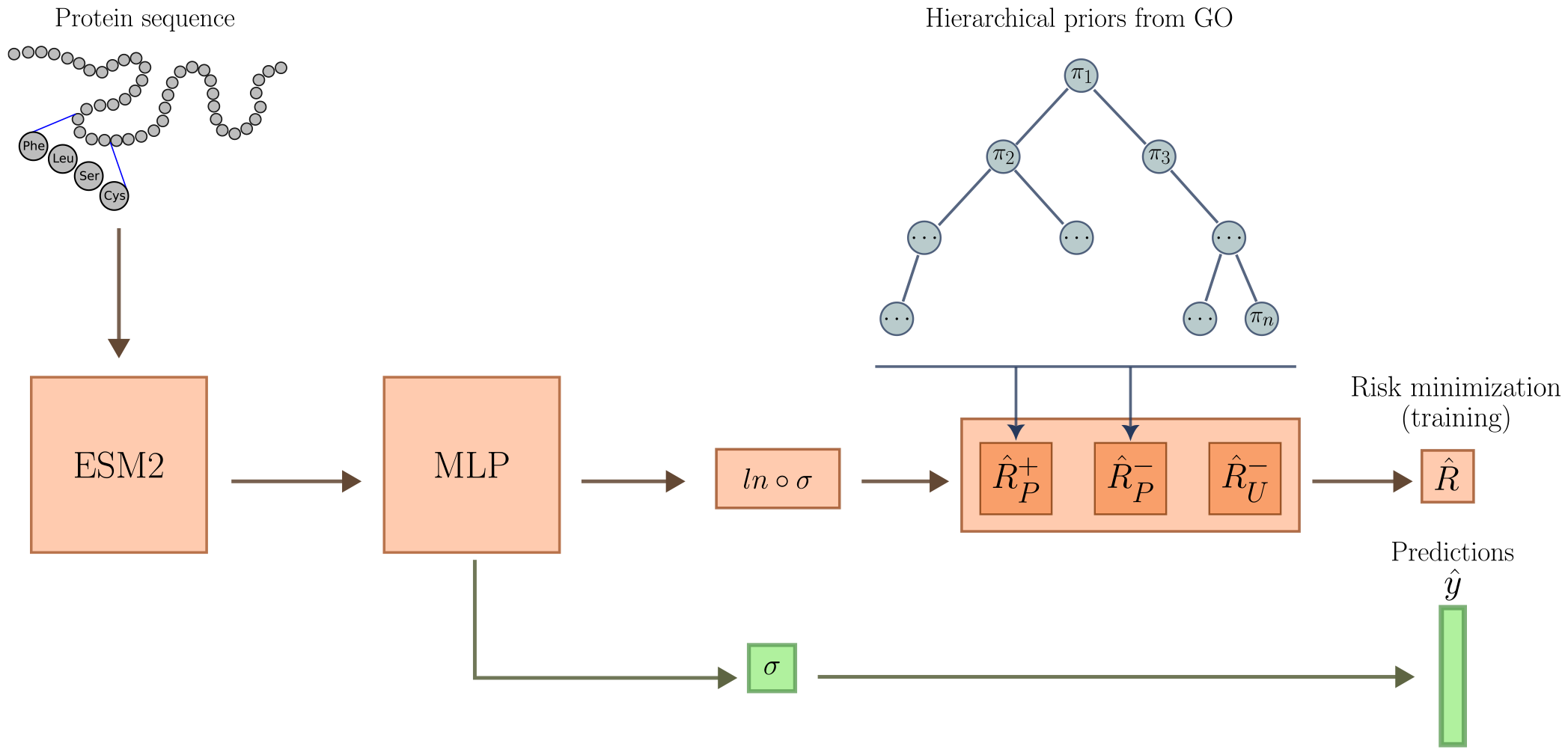
PU-GO workflow. The MLP classifier is trained to minimize classification risk of positive and unlabeled samples. Prior factors for each GO class is computed based on hierarchical GO structure.

### Multilabel PU classification

Equation 3 computes a *binary* classification risk. Function prediction of proteins is a multilabel classification problem (i.e., each protein instance can be assigned multiple functions). Thus, given *k* GO functions, the classification risk must be minimized for all the GO functions. Therefore, the classifier *g* must minimize the following risk:

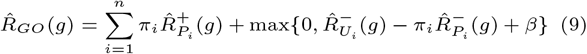

where *n* is the number of GO classes, *P*_*i*_ (*U*_*i*_) is the set of positive (unlabeled) samples for the *i*th GO function.

Additionally, the factor *π*_*i*_ = *p*(*y*_*i*_ = 1) describe the prior probability of a protein being annotated with the *i*th GO function. GO functions are structured hierarchically, which implies that all the proteins annotated to a function must also be annotated to the ontological ancestors of such function. We use this information to construct priors *π*_*i*_ in the following way: we propagate annotations from each GO function to their ancestors and compute the frequency *S*_*i*_ = *N*_*i*_*/N*_*total*_, where *N*_*i*_ is the number of training proteins annotated with the *i*th GO function and *N*_*totoal*_ is the total number of training proteins. Let *S*_*max*_ be the largest frequency, then:

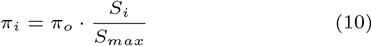

where *π*_*o*_ is a tunable hyperparameter.

### Ranking Positive and Unlabeled Samples

In Equation 9,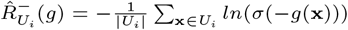. The term *σ*(−*g*(**x**)) pushed the scores to be 0, which may be unnecessarily difficult to achieve (Tang *et al*., 2022). An easier way to optimize the classifier *g* is to just push positive samples to be ranked higher than unlabeled samples. For this reason, we set:

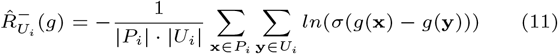

### UniProtKB/Swiss-Prot Dataset and Gene Ontology

We use the dataset that was generated from manually curated and reviewed dataset of proteins from the UniProtKB/Swiss-Prot Knowledgebase (Consortium, 2022) version 2023 03 released on 28-Jun-2023. We filtered all proteins with experimental functional annotations with evidence codes EXP, IDA, IPI, IMP, IGI, IEP, TAS, IC, HTP, HDA, HMP, HGI, HEP. The dataset contains 79, 973 reviewed and manually annotated proteins. We split this dataset into training, validation and testing sets based on sequence similarity so that there are no similar sequences to training set in the validation and testing set. We call this dataset similarity-based dataset. We use Gene Ontology (GO) released on 2023-01-01. We train and evaluate models for each of the sub-ontologies of GO separately.

To compare our model with other methods we generated a test set by following CAFA (Radivojac *et al*., 2013) challenge time-based approach. We downloaded UniProtKB/Swiss-Prot version 2023 05 released on 08-Nov-2023 and extracted newly annotated proteins in this version. Table 1 summarizes the datasets for each sub-ontology.

**Table 1.**
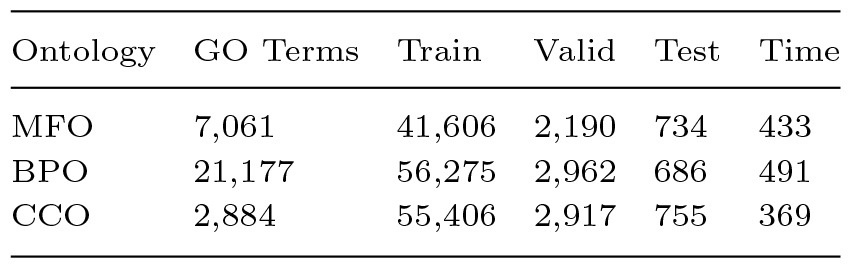
Summary of the UniProtKB/Swiss-Prot dataset.

| Ontology | GO Terms | Train | Valid | Test | Time |
| --- | --- | --- | --- | --- | --- |
| MFO | 7,061 | 41,606 | 2,190 | 734 | 433 |
| BPO | 21,177 | 56,275 | 2,962 | 686 | 491 |
| CCO | 2,884 | 55,406 | 2,917 | 755 | 369 |

### Training procedure

To train our models, we optimized hyperparameters: batch size [30, 200], margin factor [0.1, 0.01], maximum learning rate [10^−2^, 5 · 10^−6^], minimum learning rate factor [10^−1^, 10^−4^], initial prior (*π*_*o*_) [10^−3^, 10^−4^]. Hyperparameters were optimizized via Gaussian-Process Bayesian optimization method (Rasmussen and Williams, 2005; Shahriari *et al*., 2016). We used Adam (Kingma and Ba, 2015) optimizer and adapted the learning rate using a cyclic scheduler (Smith, 2017). Selected hyperparameters can be found in the Supplementary Material.

### Baseline and Comparison methods

We trained PU-GO on the similarity-based dataset in order to avoid overfitting to similar sequences. As baselines, we trained two baseline methods DeepGO-CNN (Kulmanov and Hoehndorf, 2019) and DeepGOZero (Kulmanov and Hoehndorf, 2022) and generate predictions without using any sequence similarity component such as BLAST (Altschul *et al*., 1997) or Diamond (Buchfink *et al*., 2014). For time-based dataset evaluation we selected three state-of-the-art methods such as TALE (Cao and Shen, 2021a), SPROF (Yuan *et al*., 2023b) and NetGO3 (Wang *et al*., 2023). We used baseline methods available models to generate predictions. Since baseline predictions also include sequence similarity components, we also combined PU-GO scores with Diamond predictions.

#### Naive approach

Due to the imbalance in GO class annotations and propagation based on the true-path-rule, some classes have more annotations than others. Therefore, it is possible to obtain prediction results just by assigning the same GO classes to all proteins based on annotation frequencies. In order to test the performance obtained based on annotation frequencies, CAFA introduced a baseline approach called “naive” classifier (Radivojac *et al*., 2013). Here, each query protein *p* is annotated with the GO classes with a prediction scores computed as:

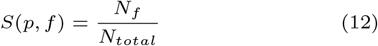

where *f* is a GO class, *N*_*f*_ is a number of training proteins annotated by GO class *f* and *N*_*total*_ is a total number of training proteins. We implement the same method.

#### MLP (ESM2)

The MLP baseline method predicts protein functions using a multi-layer perceptron (MLP) from a protein’s ESM2 embedding (Lin *et al*., 2022). We generate an embedding vector of size 5192 using ESM2 15B model and pass it to the MLP described in Equation 4. Additionally, we pass this representation to a sigmoid activation function.

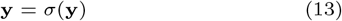

We train a different model for each sub-ontology in GO.

#### DeepGO-PLUS and DeepGOCNN

DeepGO-PLUS (Kulmanov and Hoehndorf, 2019) predicts function annotations of proteins by combining DeepGOCNN, which predicts functions from the amino acid sequence of a protein using a 1-dimensional convolutional neural network (CNN), with the DiamondScore method. DeepGOCNN captures sequence motifs that are related to GO functions. Here, we only use CNN based predictions.

#### DeepGOZero

DeepGOZero (Kulmanov and Hoehndorf, 2022) combines protein function prediction with a model-theoretic approach for embedding ontologies into a distributed geometric space. ELEmbeddings (Kulmanov *et al*., 2019) represent classes as *n*-balls and relations as vectors to embed ontology semantics into a geometric model. It uses InterPro domain annotations represented as binary vector as input and applies two layers of MLPBlock as in our MLP baseline method to generate an embedding of size 1024 for a protein. It learns the embedding space for GO classes using ELEmbeddings loss functions and optimizes together with protein function prediction loss. For a given protein *p* DeepGOZero predicts annotations for a class *c* using the following formula:

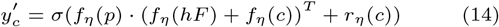

where *f*_*η*_ is an embedding function, *hF* is the <monospace> hasFunction <monospace> relation, *r*_*η*_(*c*) is the radius of an *n*-ball for a class *c* and *σ* is a sigmoid activation function. It optimizes binary crossentropy loss between predictions and the labels together with ontology axioms losses from ELEmbeddings.

#### TALE

TALE (Cao and Shen, 2021b) predicts functions using a transformer-based deep neural network model which incorporates hierarchical relations from the GO into the model’s loss function. The deep neural network predictions are combined with predictions based on sequence similarity. We used the trained models provided by the authors to evaluate them on the time-based dataset.

#### SPROF-GO

SPROF-GO (Yuan *et al*., 2023a) method uses the ProtT5-XL-U50 (Elnaggar *et al*., 2022) protein language model to extract proteins sequence embeddings and learns an attention-based neural network model. The model incorporates the hierarchical structure of GO into the neural network and predicts functions that are consistent with hierarchical relations of GO classes. Furthermore, SPROF-GO combines sequence similarity-based predictions using a homology-based label diffusion algorithm. We used the trained models provided by the authors to evaluate them on the time-based dataset.

#### NetGO3

NetGO3 integrates seven component methods that differ on the type of information they rely on: (1) Naive: GO frequency, (2) BLAST-KNN: sequence homology, (3) LR-3mer: amino acid trigram, (4) LR-InterPro: domain/family/motif, (5) NetKNN: protein network, (6) LR-Text: literature and (7) LR-ESM: protein language model. Methods with the prefix “LR” and “KNN”contain a logistic regression classifier and k-nearest neighbor algorithm, respectively. We used the web service provided by the authors to obtain predictions for our time-based benchmark dataset.

### Evaluation

We use four different measures to evaluate the performance of our models. Three protein-centric measures *F*_max_, *S*_min_ and AUPR and one class-centric AUC.

*F*_max_ is a maximum protein-centric F-measure computed over all prediction thresholds. First, we compute average precision and recall using the following formulas:

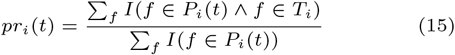

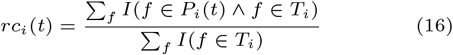

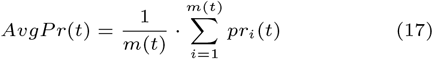

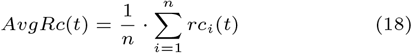

where *f* is a GO class, *T*_*i*_ is a set of true annotations, *P*_*i*_(*t*) is a set of predicted annotations for a protein *i* and threshold *t, m*(*t*) is a number of proteins for which we predict at least one class, *n* is a total number of proteins and *I* is an indicator function which returns 1 if the condition is true and 0 otherwise. Then, we compute the *F*_max_ for prediction thresholds *t* ∈ [0, 1] with a step size of 0.01. We count a class as a prediction if its prediction score is greater or equal than *t*:

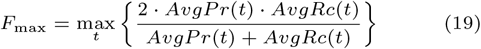

*S*_min_ computes the semantic distance between real and predicted annotations based on information content of the classes. The information content *IC*(*c*) is computed based on the annotation probability of the class *c*:

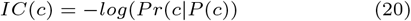

where *P* (*c*) is a set of parent classes of the class *c*. The *S*_min_ is computed using the following formulas:

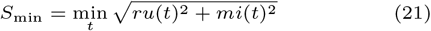

where *ru*(*t*) is the average remaining uncertainty and *mi*(*t*) is average misinformation:

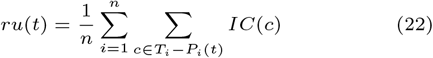

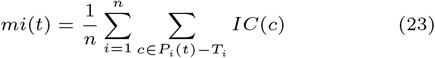

AUPR is the area under the average precision (*AvgPr*) and recall (*AvgRc*) curve. AUC is a class-centric measure where compute AUC ROC per each class and take the average.

## Results

### Prediction model: PU-GO

We developed PU-GO, a method based on positive unlabeled learning to predict GO functions. PU-GO acts on the MLP classifier shown in Equations 5–8. The training phase uses the output of the classifier to compute the classification risk of positive and unlabeled samples following Equation 9. In the prediction phase, the output of the classifier is passed to the sigmoid function directly.

We trained three separate models for each sub-ontology. The only parametric difference between the three models is the output size of the classifier, which depends of the number of GO functions. For Molecular Function Ontology there are 7,114 functions, for Cellular Component ontology 2,888 and for Biological Process Ontology 21,105.

We used the similarity-based dataset to train our models in order to avoid bias induced by sequence-similar proteins existing in training and testing datasets. For each model, we trained 10 models and the predictions are aggregated using the arithmetic mean operation.

### Evaluation on similarity-based split

To evaluate PU-GO, we chose baseline methods that do not contain components relying on sequence similarity for computing prediction scores. Results are shown in Table 2. PU-GO consistently outperforms baseline methods. While DeepGO-CNN and DeepGOZero use background knowledge to enhance protein function prediction, using unlabeled samples as negatives can incorrectly bias the model into false-negative predictions. Additionally, PU learning robustness is evidenced when comparing to MLP(ESM2), which uses the same classifier function as PU-GO but consider unlabeled samples as negatives.

**Table 2.**
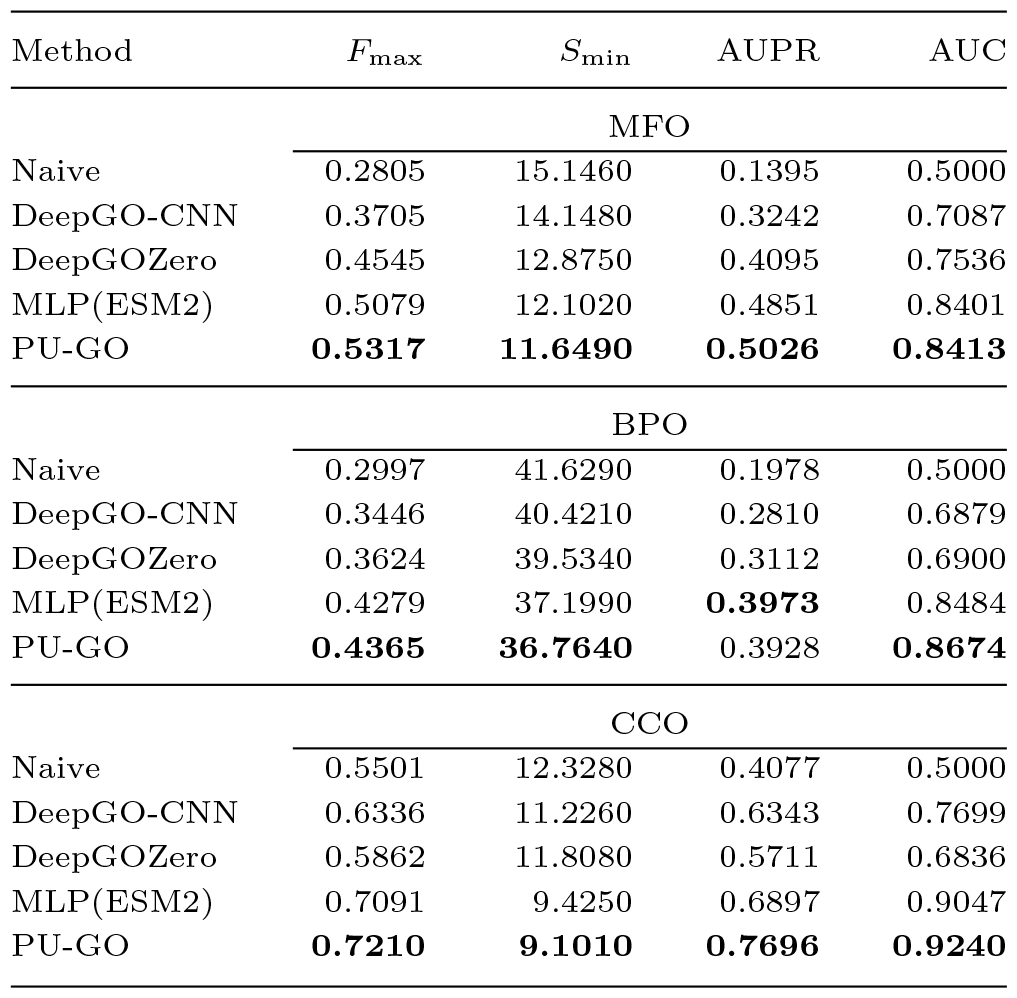
Evaluation results for similarity-based split using protein-centric *F*_max_, *S*_min_, and AUPR, and the class-centric average AUC. Method *F*_max_ *S*_min_ AUPR AUC.

### Evaluation on time-based benchmark

To test the generalization capability of PU-GO, we use our trained models, optimized using data from UniProtKB/SwissProt Knowledgebase version 2023 03, to predict GO functions from UniProtKB/SwissProt Knowledgebase version 2023 05. We compared with several state-of-the-art methods and show the results in Table 3. To enable a fair comparison with baselines, we integrate Diamond predictions to PU-GO. PU-GO+Diamond outperforms all baselines in the three subontologies. Furthermore, PU-GO alone shows comparable results to baselines, suggesting PU learning robustness to handle new annotations that other methods treat as negatives in their training.

**Table 3.**
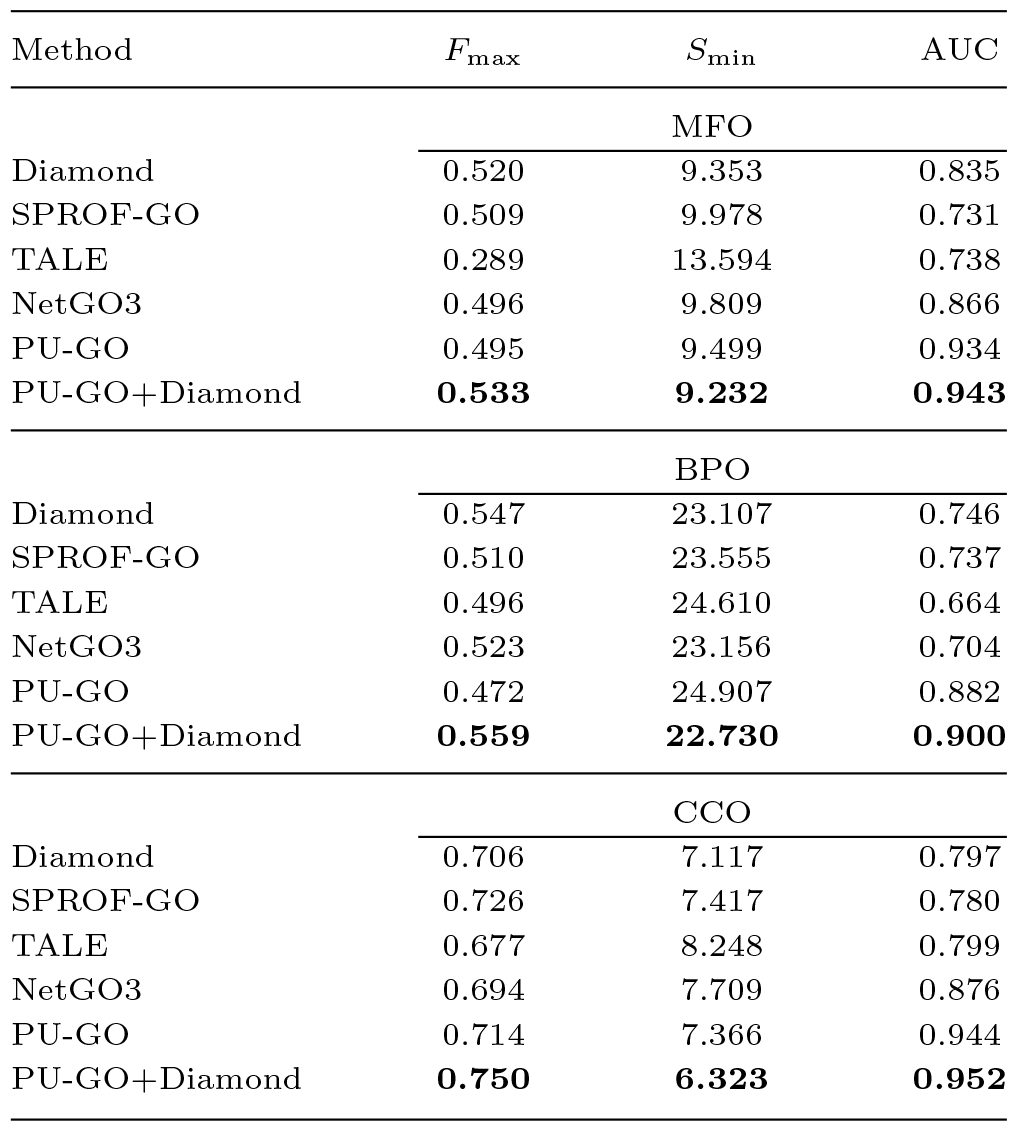
Evaluation results for time-based split using protein-centric *F*_max_, *S*_min_, and the class-centric average AUC.

### Ablation study

PU-GO contains two variations from the standard PU learning formulation such as (1) the use of a ranking loss between positive and unlabeled samples following (Tang *et al*., 2022) and (2) the use of a different prior for each GO class using GO hierarchical structure. We analyze the impact of each component in Table 4. PU-basic uses Equation 9 with *π*_*i*_ = *π*_*o*_ for every *i*th GO function. From PU-basic, we construct PU-ranking replacing the risk estimation for unlabeled samples 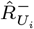 from Equation 9 with a risk computing the ranking between positive and unlabeled samples in Equation 11. PU-ranking is more flexible than PU-basic, and only requires unlabeled samples to be scored lower than positive ones and not strictly close to 0, which results in better performance in general. Finally, from PU-ranking we construct PU-GO by incorporating custom priors *π*_*i*_ for each GO class (Equation 10). This change shallowly incorporates hierarchy information as class priors (i.e, a GO class closer to the root is more likely to be annotated with a protein than a GO class closer to the leaves). Our analysis shows that using custom prior values enhance PU learning. For every method, we trained 10 models and report the mean and standard deviation values.

**Table 4.**
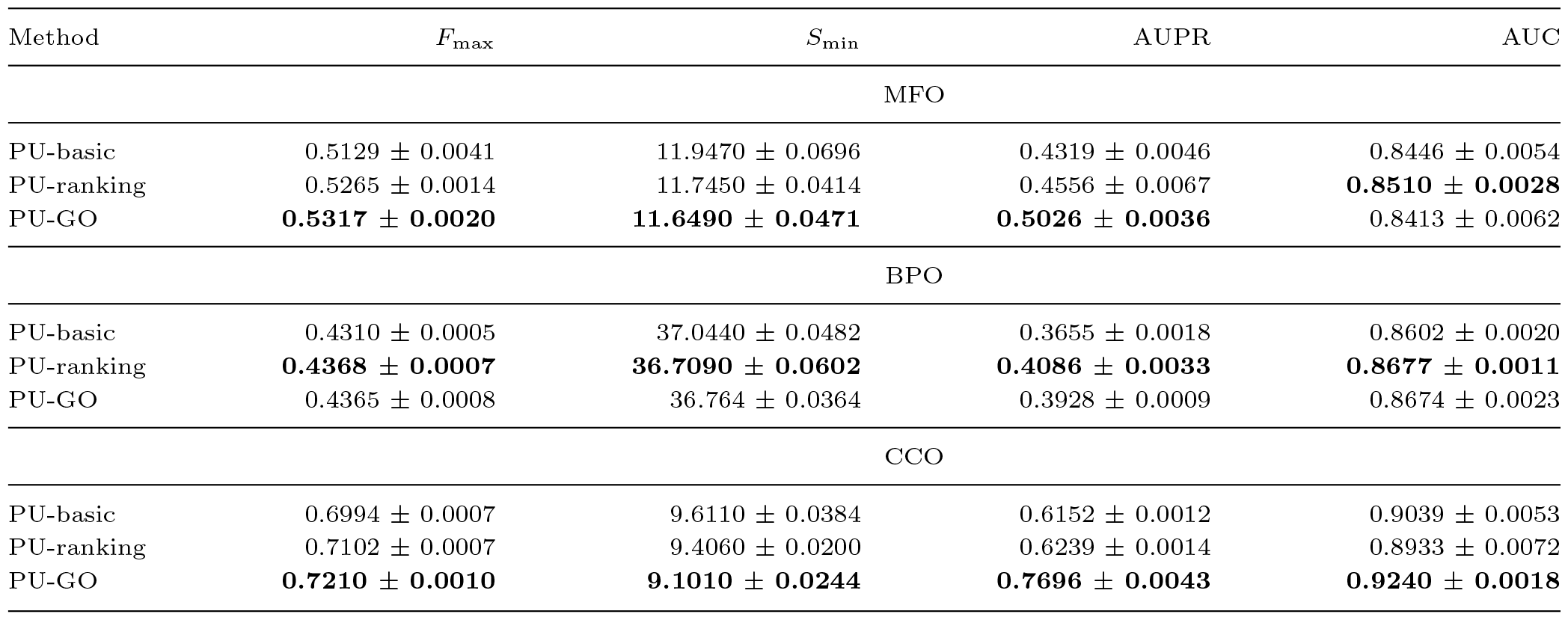
Ablation study analyzing the components of PU-GO. Metrics reported are protein-centric *F*_max_, *S*_min_, and AUPR, and the class-centric average AUC.

## Discussion

Positive-unlabeled learning is an appropriate formulation to the automated function prediction problem, where most of the data is still not labeled. Previous attempts to handle unlabeled data aim to transform some unlabeled samples into negatives (Youngs *et al*., 2013) or have not been applied to current deep learning classifiers (Song *et al*., 2021). We developed PU-GO, adapting *risk-minimization* based PU learning (Elkan and Noto, 2008; Bekker and Davis, 2020; du Plessis *et al*., 2014; Plessis *et al*., 2015; Kiryo *et al*., 2017) to the context of function prediction. PU-GO does not require extracting a subset of unlabeled samples as negatives. Instead, the whole unlabeled dataset can be used to adapt a classifier.

PU learning with risk-minimization framework is a function of a classifier. In our case, we used an MLP classifier. The input for the MLP were vectors from ESM2 15B, a pretrained language model for protein sequences. This configuration (i.e., ESM2 15B + MLP) is similar to other methods such as SPROF-GO (Yuan *et al*., 2023a), NetGO3 (Wang *et al*., 2023), which as part of their frameworks there are pretrained language models together with a classifier. PU-GO does not contain any additional component other than the ESM2 15B+MLP classifier. We showed that PU-GO was able to outperform baseline methods as well as the binary classification training version of ESM2 15B + MLP, which supports the hypothesis that PU learning is an appropriate aproach to improve protein function prediction. However, more sophisticated classifiers can be proposed in future work, where incorporation of additional domain-specific biological data can be used to constrain the optimization process.

Class prior estimation is a crucial aspect in PU learning (du Plessis *et al*., 2016). For protein function prediction, we leveraged domain-specific information such as the GO hierarchical structure to design custom class priors per each GO class based on their annotation frequency. Despite the simpicity of this approach, it showed to be effective to construct a more robust models. However, future work can explore more precise ways to construct better priors by leveraging deeper aspects of GO (not only class annotation frequency) such as semantic similarity between GO classes. Furthermore, biological information can also be leveraged to construct better class priors such as protein sequence homology (Yuan *et al*., 2023a) or protein sequence similarity.

PU-GO framework handle unlabeled samples differently than previous approaches where the aim was to strictly discriminate between positive and negative samples. In PU-GO, instead of minimizing the risk of clasifying an unlabeled sample as negative, it adresses the protein function prediction as a ranking problem and minimizes the risk of ranking an unlabeled sample higher than a positive one. Furthermore, since the risk-minimization framework we resort to is extensible to incorporate true negative samples (Hsieh *et al*., 2019), future work can be directed to study the incorporation of negative annotations that are already available or that can be extracted by some strategy.

## Conclusion

Protein function prediction is a widely studied multilabel classification problem that typically has been addressed under binary classification settings. However, protein function annotations are mostly unlabeled. To deal with unlabeled annotations, we addressed protein function prediction as a PU classification problem. We adapted the PU learning framework for protein function prediction by incorporating hierarchical information in GO in the class priors. Our analysis indicates improved performance compared to existing methods on similarity-based and time-based benchmark datasets. Future potential work could focus on incorporating negative samples to the PU setting and minimize negative classification risk. Although negative data is small, finding a way to use it can improve the classifier generalization capability. Another direction could be using more sophisticated classifiers that can include other types of biological information, which has been an approach followed in the binary-classification setting.

## Author contributions statement

R.H., M.K., Z.T. and F.Z. conceived the experiment(s), M.K. and F.Z. conducted the experiment(s), R.H., M.K. and F.Z. analysed the results. R.H., M.K., Z.T. and F.Z. wrote and reviewed the manuscript. R.H. supervised the work. R.H. and M.K. acquired funding.

## Acknowledgments

This work has been supported by funding from King Abdullah University of Science and Technology (KAUST) Office of Sponsored Research (OSR) under Award No. URF/1/4355-01-01, URF/1/4675-01-01, URF/1/4697-01-01, URF/1/5041-01-01, REI/1/5659-01-01, and FCC/1/1976-46-01. This work was supported by the SDAIA-KAUST Center of Excellence in Data Science and Artificial Intelligence (SDAIA-KAUST AI). We acknowledge support from the KAUST Supercomputing Laboratory.

## References

Altschul, S. F., Madden, T. L., Schäffer, A. A., Zhang, J., Zhang, Z., Miller, W., and Lipman, D. J. (1997). Gapped BLAST and PSI-BLAST: a new generation of protein database search programs. Nucleic Acids Research, 25(17), 3389–3402.

Ashburner, M., Ball, C. A., Blake, J. A., Botstein, D., Butler, H., Cherry, M. J., Davis, A. P., Dolinski, K., Dwight, S. S., Eppig, J. T., Harris, M. A., Hill, D. P., Tarver, L. I., Kasarskis, A., Lewis, S., Matese, J. C., Richardson, J. E., Ringwald, M., Rubin, G. M., and Sherlock, G. (2000). Gene ontology: tool for the unification of biology. Nature Genetics, 25(1), 25–29.

Bekker, J. and Davis, J. (2020). Learning from positive and unlabeled data: a survey. Machine Learning, 109(4), 719–760.

Buchfink, B., Xie, C., and Huson, D. H. (2014). Fast and sensitive protein alignment using diamond. Nature Methods, 12, 59 EP –. [PubMed:25402007] [doi:10.1038/nmeth.3176].

Cao, Y. and Shen, Y. (2021a). TALE: Transformer-based protein function Annotation with joint sequence–Label Embedding. Bioinformatics, 37(18), 2825–2833.

Cao, Y. and Shen, Y. (2021b). TALE: Transformer-based protein function Annotation with joint sequence–Label Embedding. Bioinformatics, 37(18), 2825–2833.

Chen, Y., Li, Z., Wang, X., Feng, J., and Hu, X. (2010). Predicting gene function using few positive examples and unlabeled ones. BMC Genomics, 11(Suppl 2), S11.

Consortium, T. U. (2022). UniProt: the Universal Protein Knowledgebase in 2023. Nucleic Acids Research, 51(D1), D523–D531.

Cortes, C. and Vapnik, V. (1995). Support-vector networks. Machine Learning, 20(3), 273–297.

du Plessis, M. C., Niu, G., and Sugiyama, M. (2014). Analysis of learning from positive and unlabeled data. In Z. Ghahramani, M. Welling, C. Cortes, N. Lawrence, and K. Weinberger, editors, Advances in Neural Information Processing Systems, volume 27. Curran Associates, Inc.

du Plessis, M. C., Niu, G., and Sugiyama, M. (2016). Class-prior estimation for learning from positive and unlabeled data. Machine Learning, 106(4), 463–492.

Eisenberg, D., Marcotte, E. M., Xenarios, I., and Yeates, T. O. (2000). Protein function in the post-genomic era. Nature, 405(6788), 823–826.

Elkan, C. and Noto, K. (2008). Learning classifiers from only positive and unlabeled data. In Proceedings of the 14th ACM SIGKDD International Conference on Knowledge Discovery and Data Mining, KDD ‘08, page 213–220, New York, NY, USA. Association for Computing Machinery.

Elnaggar, A., Heinzinger, M., Dallago, C., Rehawi, G., Wang, Y., Jones, L., Gibbs, T., Feher, T., Angerer, C., Steinegger, M., Bhowmik, D., and Rost, B. (2022). Prottrans: Toward understanding the language of life through self-supervised learning. IEEE Transactions on Pattern Analysis and Machine Intelligence, 44(10), 7112–7127.

Hsieh, Y.-G., Niu, G., and Sugiyama, M. (2019). Classification from positive, unlabeled and biased negative data. In K. Chaudhuri and R. Salakhutdinov, editors, Proceedings of the 36th International Conference on Machine Learning, volume 97 of Proceedings of Machine Learning Research, pages 2820–2829. PMLR.

Kenneth Lange, D. R. H. and Yang, I. (2000). Optimization transfer using surrogate objective functions. Journal of Computational and Graphical Statistics, 9(1), 1–20.

Kingma, D. P. and Ba, J. (2015). Adam: A method for stochastic optimization. In Y. Bengio and Y. LeCun, editors, 3rd International Conference on Learning Representations, ICLR 2015, San Diego, CA, USA, May 7–9, 2015, Conference Track Proceedings.

Kiryo, R., Niu, G., du Plessis, M. C., and Sugiyama, M. (2017). Positive-unlabeled learning with non-negative risk estimator. In I. Guyon, U. V. Luxburg, S. Bengio, H. Wallach, R. Fergus, S. Vishwanathan, and R. Garnett, editors, Advances in Neural Information Processing Systems, volume 30. Curran Associates, Inc.

Kulmanov, M. and Hoehndorf, R. (2019). DeepGOPlus: improved protein function prediction from sequence. Bioinformatics. [PubMed:31350877] [doi:10.1093/bioinformatics/btz595].

Kulmanov, M. and Hoehndorf, R. (2022). DeepGOZero: improving protein function prediction from sequence and zero-shot learning based on ontology axioms. Bioinformatics, 38(Supplement 1), i238–i245.

Kulmanov, M., Khan, M. A., and Hoehndorf, R. (2017). DeepGO: predicting protein functions from sequence and interactions using a deep ontology-aware classifier. Bioinformatics, 34(4), 660–668. [PubMed:29028931] [PubMed Central:PMC5860606] [doi:10.1093/bioinformatics/btx624].

Kulmanov, M., Liu-Wei, W., Yan, Y., and Hoehndorf, R. (2019). El embeddings: Geometric construction of models for the description logic el++. In Proceedings of the Twenty-Eighth International Joint Conference on Artificial Intelligence, IJCAI-19, pages 6103–6109. International Joint Conferences on Artificial Intelligence Organization.

Lan, W., Wang, J., Li, M., Liu, J., Li, Y., Wu, F.-X., and Pan, Y. (2016). Predicting drug–target interaction using positive-unlabeled learning. Neurocomputing, 206, 50–57. SI:DMSB.

Li, F., Dong, S., Leier, A., Han, M., Guo, X., Xu, J., Wang, X., Pan, S., Jia, C., Zhang, Y., Webb, G. I., Coin, L. J. M., Li, C., and Song, J. (2021). Positive-unlabeled learning in bioinformatics and computational biology: a brief review. Briefings in Bioinformatics, 23(1).

Lin, Z., Akin, H., Rao, R., Hie, B., Zhu, Z., Lu, W., Smetanin, N., dos Santos Costa, A., Fazel-Zarandi, M., Sercu, T., Candido, S., et al. (2022). Language models of protein sequences at the scale of evolution enable accurate structure prediction. bioRxiv .

Liu, W., Wu, A., Pellegrini, M., and Wang, X. (2015). Integrative analysis of human protein, function and disease networks. Scientific Reports, 5(1).

Peng, L., Zhu, W., Liao, B., Duan, Y., Chen, M., Chen, Y., and Yang, J. (2017). Screening drug-target interactions with positive-unlabeled learning. Scientific Reports, 7(1).

Plessis, M. D., Niu, G., and Sugiyama, M. (2015). Convex formulation for learning from positive and unlabeled data. In F. Bach and D. Blei, editors, Proceedings of the 32nd International Conference on Machine Learning, volume 37 of Proceedings of Machine Learning Research, pages 1386–1394, Lille, France. PMLR.

Radivojac, P., Clark, W. T., Oron, T. R., Schnoes, A. M., Wittkop, T., Sokolov, A., Graim, K., Funk, C., Verspoor, K., Ben-Hur, A., Pandey, G., Yunes, J. M., Talwalkar, A. S., Repo, S., Souza, M. L., Piovesan, D., Casadio, R., Wang, Z., Cheng, J., Fang, H., Gough, J., Koskinen, P., Toronen, P., Nokso-Koivisto, J., Holm, L., Cozzetto, D., Buchan, D. W. A., Bryson, K., Jones, D. T., Limaye, B., Inamdar, H., Datta, A., Manjari, S. K., Joshi, R., Chitale, M., Kihara, D., Lisewski, A. M., Erdin, S., Venner, E., Lichtarge, O., Rentzsch, R., Yang, H., Romero, A. E., Bhat, P., Paccanaro, A., Hamp, T., Kaszner, R., Seemayer, S., Vicedo, E., Schaefer, C., Achten, D., Auer, F., Boehm, A., Braun, T., Hecht, M., Heron, M., Honigschmid, P., Hopf, T. A., Kaufmann, S., Kiening, M., Krompass, D., Landerer, C., Mahlich, Y., Roos, M., Bjorne, J., Salakoski, T., Wong, A., Shatkay, H., Gatzmann, F., Sommer, I., Wass, M. N., Sternberg, M. J. E., Skunca, N., Supek, F., Bosnjak, M., Panov, P., Dzeroski, S., Smuc, T., Kourmpetis, Y. A. I., van Dijk, A. D. J., Braak, C. J. F. t., Zhou, Y., Gong, Q., Dong, X., Tian, W., Falda, M., Fontana, P., Lavezzo, E., Di Camillo, B., Toppo, S., Lan, L., Djuric, N., Guo, Y., Vucetic, S., Bairoch, A., Linial, M., Babbitt, P. C., Brenner, S. E., Orengo, C., Rost, B., Mooney, S. D., and Friedberg, I. (2013). A large-scale evaluation of computational protein function prediction. Nat Meth, 10(3), 221–227. [PubMed:23353650] [PubMed Central:PMC3584181] [doi:10.1038/nmeth.2340].

Rasmussen, C. E. and Williams, C. K. I. (2005). Gaussian processes for machine learning. Adaptive Computation and Machine Learning series. MIT Press, London, England.

Schenone, M., Dančík, V., Wagner, B. K., and Clemons, P. A. (2013). Target identification and mechanism of action in chemical biology and drug discovery. Nature Chemical Biology, 9(4), 232–240.

Shahriari, B., Swersky, K., Wang, Z., Adams, R. P., and de Freitas, N. (2016). Taking the human out of the loop: A review of bayesian optimization. Proceedings of the IEEE, 104(1), 148–175.

Smith, L. N. (2017). Cyclical learning rates for training neural networks. In 2017 IEEE Winter Conference on Applications of Computer Vision (WACV), pages 464–472.

Song, H., Bremer, B. J., Hinds, E. C., Raskutti, G., and Romero, P. A. (2021). Inferring protein sequence-function relationships with large-scale positive-unlabeled learning. Cell Systems, 12(1), 92–101.e8.

Stolfi, P., Mastropietro, A., Pasculli, G., Tieri, P., and Vergni, D. (2023). NIAPU: network-informed adaptive positive-unlabeled learning for disease gene identification. Bioinformatics, 39(2), btac848.

Tang, Z., Pei, S., Zhang, Z., Zhu, Y., Zhuang, F., Hoehndorf, R., and Zhang, X. (2022). Positive-unlabeled learning with adversarial data augmentation for knowledge graph completion. In L. D. Raedt, editor, Proceedings of the Thirty-First International Joint Conference on Artificial Intelligence, IJCAI-22, pages 2248–2254. International Joint Conferences on Artificial Intelligence Organization. Main Track.

Vasighizaker, A. and Jalili, S. (2018). C-pugp: A cluster-based positive unlabeled learning method for disease gene prediction and prioritization. Computational Biology and Chemistry, 76, 23–31.

Wang, S., You, R., Liu, Y., Xiong, Y., and Zhu, S. (2023). Netgo 3.0: Protein language model improves large-scale functional annotations. Genomics, Proteomics & Bioinformatics, 21(2), 349–358.

Yang, P., Li, X.-L., Mei, J.-P., Kwoh, C.-K., and Ng, S.-K. (2012). Positive-unlabeled learning for disease gene identification. Bioinformatics, 28(20), 2640–2647.

Youngs, N., Penfold-Brown, D., Drew, K., Shasha, D., and Bonneau, R. (2013). Parametric bayesian priors and better choice of negative examples improve protein function prediction. Bioinformatics, 29(9), 1190–1198.

Yuan, Q., Xie, J., Xie, J., Zhao, H., and Yang, Y. (2023a). Fast and accurate protein function prediction from sequence through pretrained language model and homology-based label diffusion. Briefings in Bioinformatics, 24(3), bbad117.

Yuan, Q., Xie, J., Xie, J., Zhao, H., and Yang, Y. (2023b). Fast and accurate protein function prediction from sequence through pretrained language model and homology-based label diffusion. Briefings in Bioinformatics.

Zhao, X.-M., Wang, Y., Chen, L., and Aihara, K. (2008). Gene function prediction using labeled and unlabeled data. BMC Bioinformatics, 9(1).

